# Expanding the synthetic biology toolbox with a library of constitutive and repressible promoters

**DOI:** 10.1101/2022.10.10.511673

**Authors:** Eric J.Y. Yang, Jennifer L. Nemhauser

## Abstract

**Background:** To support the increasingly complex circuits needed for plant synthetic biology applications, additional constitutive promoters are essential. Reusing promoter parts can lead to difficulty in cloning, increased heterogeneity between transformants, transgene silencing and trait instability. Moreover, the utility of such promoters could be increased by introducing target sequences not found elsewhere in the *Arabidopsis thaliana* genome and well-suited for Cas9-associated guide RNAs (gRNAs).

**Methods:** We have developed a pipeline to identify genes that have stable expression across a wide range of *Arabidopsis* tissues at different developmental stages, and have identified a number of promoters that are well expressed in both transient (*Nicotiana benthamiana*) and stable (*Arabidopsis*) transformation assays. We have also introduced two genome-orthogonal gRNA target-sites in a subset of the screened promoters, converting them into NOR logic gates.

**Results:** Of twenty-two promoters identified in our bioinformatic screen, sixteen drove detectable reporter expression in *N. benthamiana*. Only three of these promoters were able to produce visible expression of the RUBY reporter in *Arabidopsis* despite producing RUBY mRNA that could be readily detected by qPCR. We then modified six of these promoters to be repressible, and five of which functioned as NOR gates

**Conclusions:** One of the major bottlenecks for the ambitious engineering projects currently under development in plants is the lack of well-characterized constitutive promoters. The work here begins to fill this gap. It can also form the basis of constructing more complex information processing circuits in the future.

## Introduction

Plant synthetic biology aims to provide greater control over plant form and function, a goal that is beginning to be realized. Several projects have produced measurable gains in photosynthetic efficiency [1,2], and others have intervened in hormone response pathways to change plant architecture [3] or environmental response [4,5]. These advances rely on well-characterized promoters to ensure expression of transgene in desired tissues.

Promoters can be broadly broken down into three categories based on expression pattern: constitutive, spatiotemporally-restricted, and inducible [6]. Constitutive promoters are expressed in all tissues at all times, and regulate the transcription of what are commonly referred to as “housekeeping genes”. While each category of promoter is useful in plant engineering, constitutive promoters are often used to confer novel traits such as herbicide tolerance, to drive synthetic circuits, and used in metabolic engineering projects due to their broad tissue coverage [7–9]. Some of the most widely used plant constitutive promoters include variants from the *Cauliflower Mosaic Virus* 35S (35S) promoter, and promoters from members of the ubiquitin and actin families [6,10]. However, the list of available plant constitutive promoters is short, and this lack of parts poses many challenges [6]. Having to reuse the limited number of promoters in increasingly complex plant gene circuits or metabolic engineering projects can quickly lead to instability of the transformed construct due to repeated elements rearranging and homology-dependent gene silencing, which is heritable [6,11,12].

Recently, several groups have used distinct strategies to overcome this engineering obstacle. One approach builds synthetic promoters by adding cis-elements to a “minimal promoter region”, which is often 35S-derived. By varying the number and type of cis-elements, researchers were able to generate promoters with a wide range of expression levels and expression patterns [8,13–15]. Another approach uses sequences upstream of the minimal promoter region as a landing dock for synthetic activators guided by zinc-finger, TALE, or dCas9 to promote expression [14]. The expression strength of these promoters can be tuned by varying the number of target sites for the synthetic activators [16,17]. These approaches, while quite powerful, are limited by the small number of characterized minimal promoters available to build upon and may still lead to repeated units in large constructs if the same minimal promoters were used.

Here, we employed an alternative approach for finding constitutive promoters. Instead of building and testing synthetic promoters, we looked to natural promoters found in the *Arabidopsis* genome. This approach has a few advantages. Synthetic promoters require extensive characterization to determine their expression pattern and, because of practical constraints, are often only tested in a few selected tissues. In contrast, the wealth of RNA-seq data available for *Arabidopsis* provides highly detailed information about a given promoter’s likely expression potential, including the expression level of the gene throughout many developmental stages, tissue types, and even various growth and/or stress conditions. The expression of a native promoter has already been subject to selective pressures, and so is potentially more likely to remain stable across generations. By introducing a set of unique sequences, natural promoters also have the potential to minimize the likelihood of gene silencing or unwanted recombination through repeated units in multigenic constructs. Lastly, by employing some of the techniques in generating synthetic promoters described above, these native promoters could potentially form the foundation for suites of derived promoters with even more refined expression levels. A similar approach successfully expanded the range of promoter expressions available in *B. subtilis* [18]. Since the argument for the need of additional promoter parts can be directly extended to the need for additional terminators, and terminators are known regulators of gene expression [19], we screened promoter-terminator pairs together. To further extend the utility of the new promoter/terminator pairs, we also introduced dCas9 target-sites with sequences not found elsewhere in the *Arabidopsis* genome, thereby enabling specific repression by synthetic transcription factors without interfering with the cognate native genes.

## Results

To identify the most stably expressing promoters available in the *Arabidopsis* genome, we analyzed publicly available RNAseq datasets. The majority of the RNAseq dataset came from the Klepikova transcriptome profile which included multiple tissues from different development stages [20]. We supplemented this dataset with an RNAseq dataset for pollen [21], as this cell-type was not represented in the Klepikova dataset. After processing the RNAseq datasets, there were 10,096 genes that were expressed in all the datasets (i.e. have none zero read counts) (Figure1A). Coefficient of variation (CV) of expression across different tissues is often used as a metric for identifying stably expressed genes [22–24]. Within the lowest 3% CV, there were 303 genes, which corresponds to a CV cutoff of 0.26 (Figure1A,D). To facilitate dissemination of the parts quantified in this study, we adopted the Golden Gate MoClo system and cloned the promoter + 5’UTR together as a standard MoClo part and similarly with 3’UTR + terminator (Figure1C) [25,26]. Since MoClo uses BsaI and BbsI type-II restriction enzymes for cloning, we removed any candidates with their corresponding restriction sites within the cloned regions. These cloning constraints left us with 61 candidate genes.

**Figure1.**
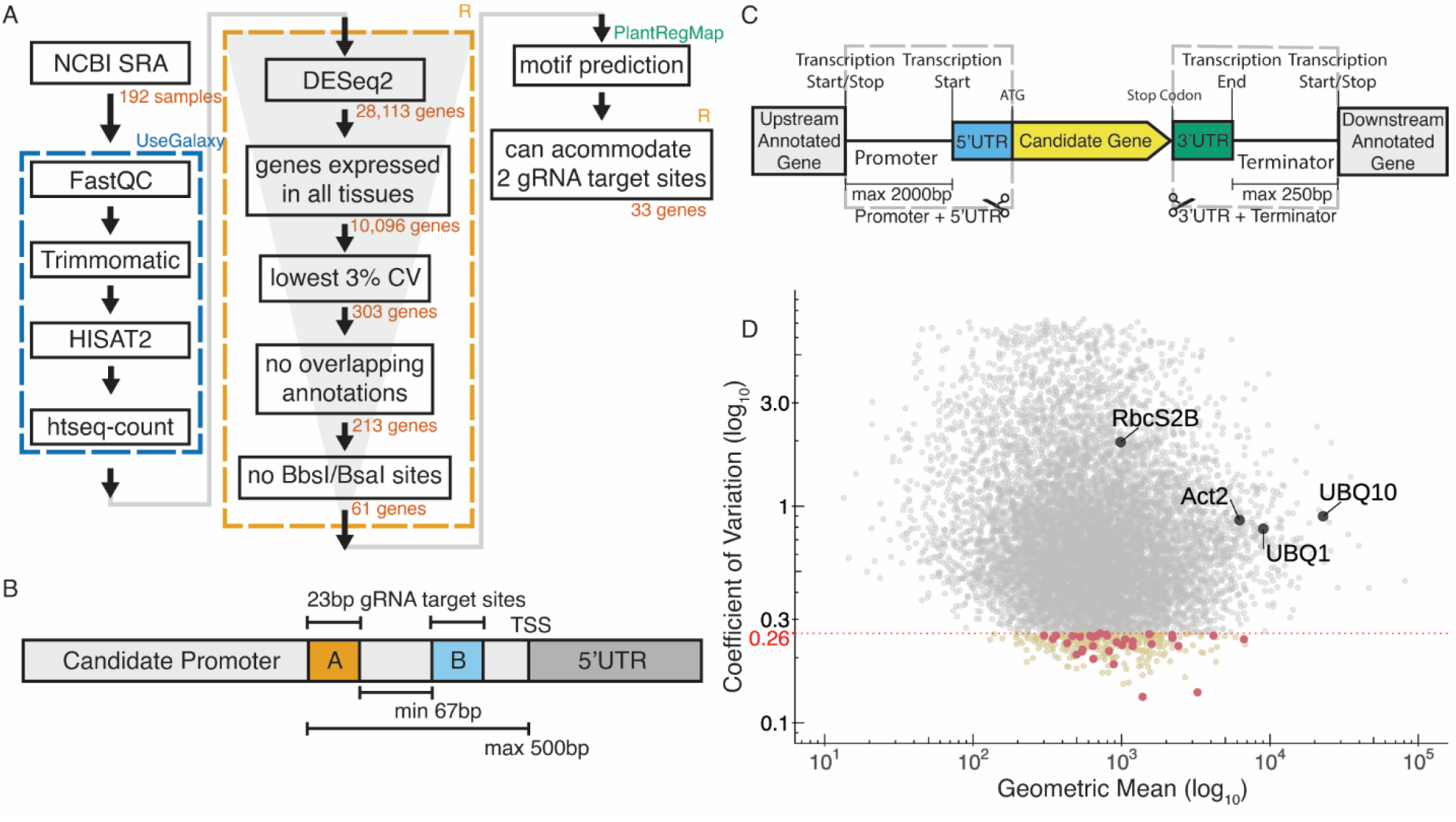
A) Pipeline to identify constitutive promoters. The number of genes that pass each filter are indicated, along with the software used to implement the analysis. Detailed methods, including parameters for each filter, are described in the Methods section. B) Schematic of filters used to select candidate promoters to engineer with synthetic gRNA target-sites. C) Schematic describing how we defined “promoter” and “terminator”. The “promoter” was defined here as starting from the transcription start site and going upstream to a maximum of 2000bp or to the next annotated neighboring gene, whichever is shorter. Similarly, a “terminator” was defined as starting from the transcription end site and going downstream a maximum of 250bp or to the next annotated neighboring gene, whichever is shorter. Promoters and terminators were cloned, along with their respective UTRs, following the Golden Gate MoClo system. D) Plot showing values for the 10,096 genes expressed in all tissues. The geometric mean of expression across samples is plotted on the x-axis with the coefficient of variation (CV) on the y-axis. Both axes are on a base-10 log scale. Lowest 3% CV corresponds to a 0.26 CV cutoff, and the 303 genes with CV lower than 0.26 are highlighted yellow. The final 33 candidates that fulfilled all criteria are highlighted in red. Several common promoters used in plant synthetic biology are annotated for reference.

To selectively activate or repress promoters in the context of a synthetic circuit, we wanted to modify segments of the promoter region to allow genome-orthogonal dCas9 targeting. Using the “Binding Site Prediction” function from PlantRegMap we screened for predicted motifs within 500 bp of the promoter region from the TSS [27]. We retained promoters that could accommodate two 23bp gRNA target-sites (20bp target sequence and 3bp PAM site) without interrupting any predicted motifs and were at least 67bp apart, following the spacing used in Gander et al. (Figure1B). We were left with 33 candidate genes. Compared to the commonly used native *Arabidopsis* constitutive promoters, the candidates identified here were more stably expressed but have mostly weaker mean expression (Figure1D). Detailed information of the 33 candidates can be found in Supplemental Table2.

While one of the main goals of this study is to identify the best available natural stable genes through analysis of RNAseq data, the “stability” of the candidate genes we screened for in this paper is constrained by the choice of RNAseq dataset used. The Klepikova dataset included stress-treated leaf samples with heat, cold, and wounding treatments, but they were not included in the CV calculation since the samples were only collected from mature third leaves and no other tissue types. Instead, we normalized the stress data with untreated “mature whole third leaf” and calculated their CV and included the result in Supplemental Table2 for reference. Similarly, while the datasets capture coarse temporal resolutions throughout development, they cannot identify the fluctuation of circadian genes and therefore we supplemented the final table with identified circadian genes from CGDB for reference [28].

Of the 33 stably expressed genes identified from the bioinformatics pipeline, we successfully cloned twenty-two promoter-terminator pairs. We tested the promoters in *Nicotiana benthamiana* (tobacco) transient agroinfiltration assays and identified sixteen promoters that had expression that are significantly different from the negative control (Figure2).

**Figure2.**
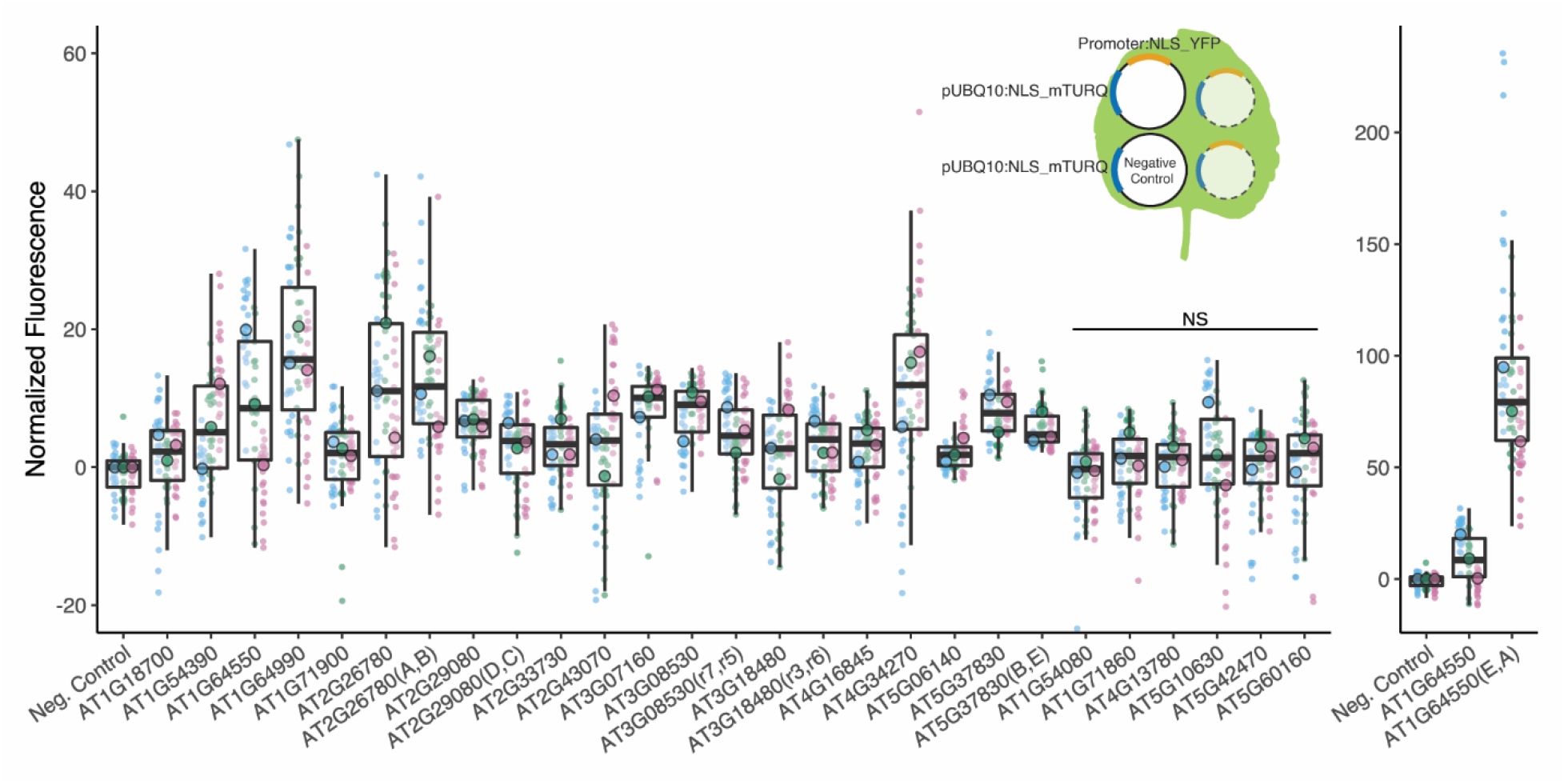
We identified sixteen promoters that expressed in *N. benthamiana*. Six promoters were modified to introduce gRNA target sites. These sites are designated by brackets following the gene name. Three different constructs were injected per leaf, each containing a promoter to be tested driving NLS_YFP and an internal control of pUBQ10:NLS_mTURQ. Each leaf also has an negative control injection that only contains pUBQ10:NLS_mTURQ. Normalization is performed using the formula: 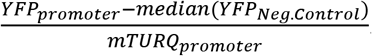. Each biological replicate is represented by a beeswarm plot of individual datapoints collected from the plate reader as well as a single summarizing datapoint representing the median. The boxplot represents all replicates. Significant test was performed using Dunnett’s test for comparing multiple treatments with control at 95% family-wise confidence level. Non-significant constructs are marked as NS. For a given construct, the colors signify datapoints derived from the same biological replicate.

To determine whether the promoters showed constitutive expression in *Arabidopsis*, twelve of the promoters were selected to drive expression of the RUBY reporter [29] in stable transformants. Since RUBY is a pigment that allows for simple visual readout [29], we were hoping it would be an effective way of evaluating the expression of the promoters in all the tissues throughout development. Three representative T1 lines were selected for each construct and six T2s per T1 line were observed at the seedling stage (12 days) and as mature plants (day 34). Eleven of the twelve promoters transformed showed expression in *N. benthamiana*, yet we only identified three promoters that displayed RUBY expression in *Arabidopsis*. Representative individuals are shown in Figure3A (Supplement Figure1).

**Figure3.**
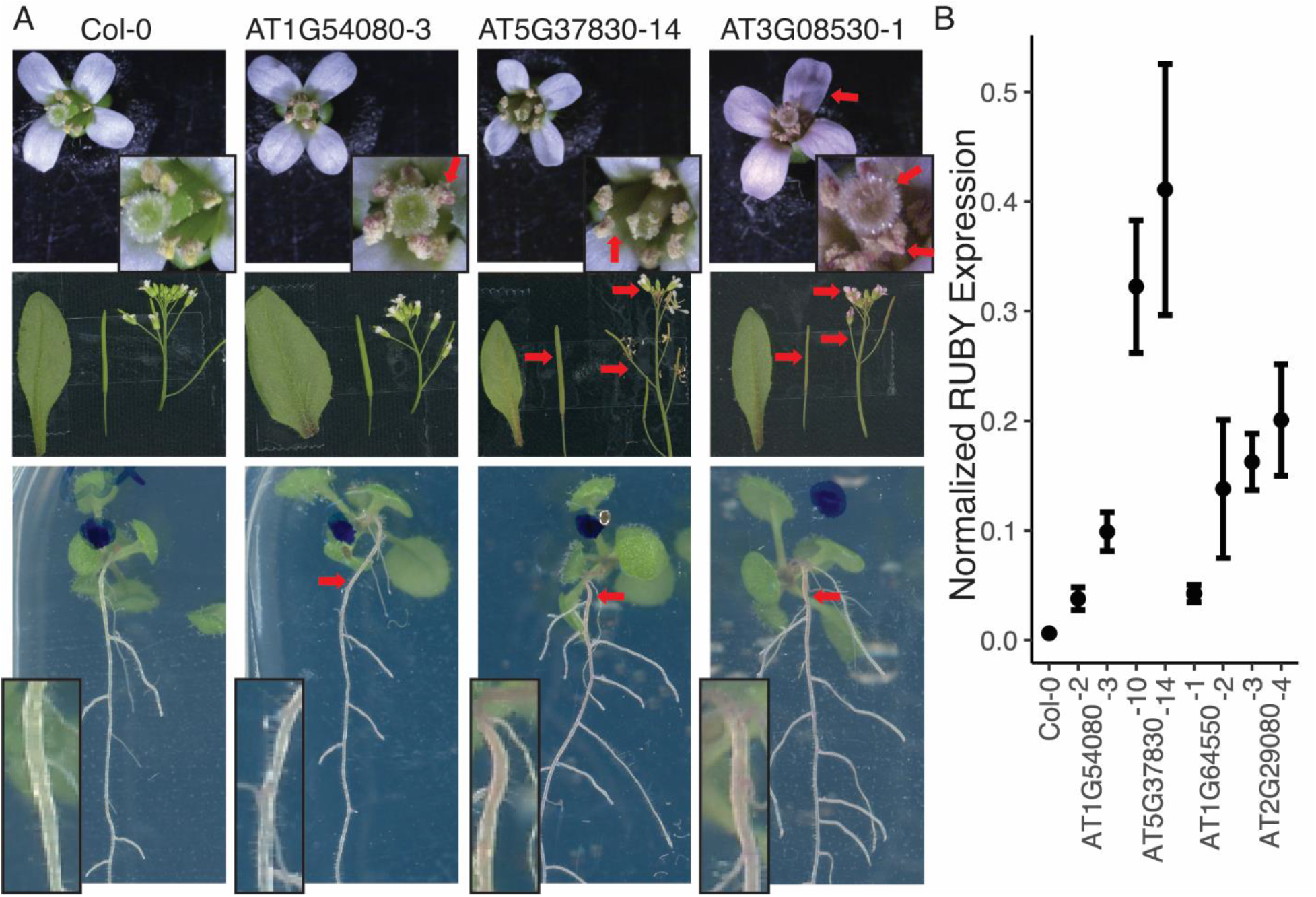
A) Three promoters showed expression of RUBY in *Arabidopsis* T2 plants. The flowers, siliques, and leaves were imaged on day 34, while the seedling images were imaged on day 12. The inset boxes are the same images at higher magnification. Red arrows indicate areas where RUBY expression is visible by eye. B) qPCR data on T2 seedlings, showing the mean expression of the RUBY reporter in each line and the standard error of the mean (SEM).

AT1G54080 displayed RUBY expression in roots and pollen. AT5G37830 had visible expression in pollen, siliques, stems, and roots. AT3G08530 had the most ubiquitous expression and had visible expression in the flowers, pollen, siliques, stems, and roots. A visual summary of the *Arabidopsis* and *N. benthamiana* experiments can be found in Supplement Figure2.

Given that the majority of the promoters had no visible expression of RUBY by eye, we performed qPCR on whole seedlings from four promoter lines: two with visible RUBY expression in roots (AT1G54080 and AT5G37830) and two without (AT1G64550 and AT2G29080). The two lines without visible RUBY expression both had qPCR expression level between the two lines with visible RUBY expression in the roots. This result suggests that RUBY was not a reliable reporter for these low expressing promoters, and that the promoters were indeed functional in *Arabidopsis* (Figure3B).

To make the constitutive promoters screened in this experiment more versatile, we next introduced two gRNA target-sites into six of the promoters screened with target-site sequences not found in the *Arabidopsis* genome. A constitutive promoter with two unique gRNA target-sites can function as a NOR gate (a two-input logic gate where the output is only ON when neither of the inputs are present) in the presence of a dCas9-guided repressor. The inputs for such a gate are the gRNAs. When either or both of the gRNAs are present, the dCas9-guided repressor should be able to keep the promoter OFF (Figure4B). Only when neither of the guides are present can the promoter be turned ON. Nine different functional gRNA sequences (A-I) were selected from the literature (Supplement Table 3).

**Figure4.**
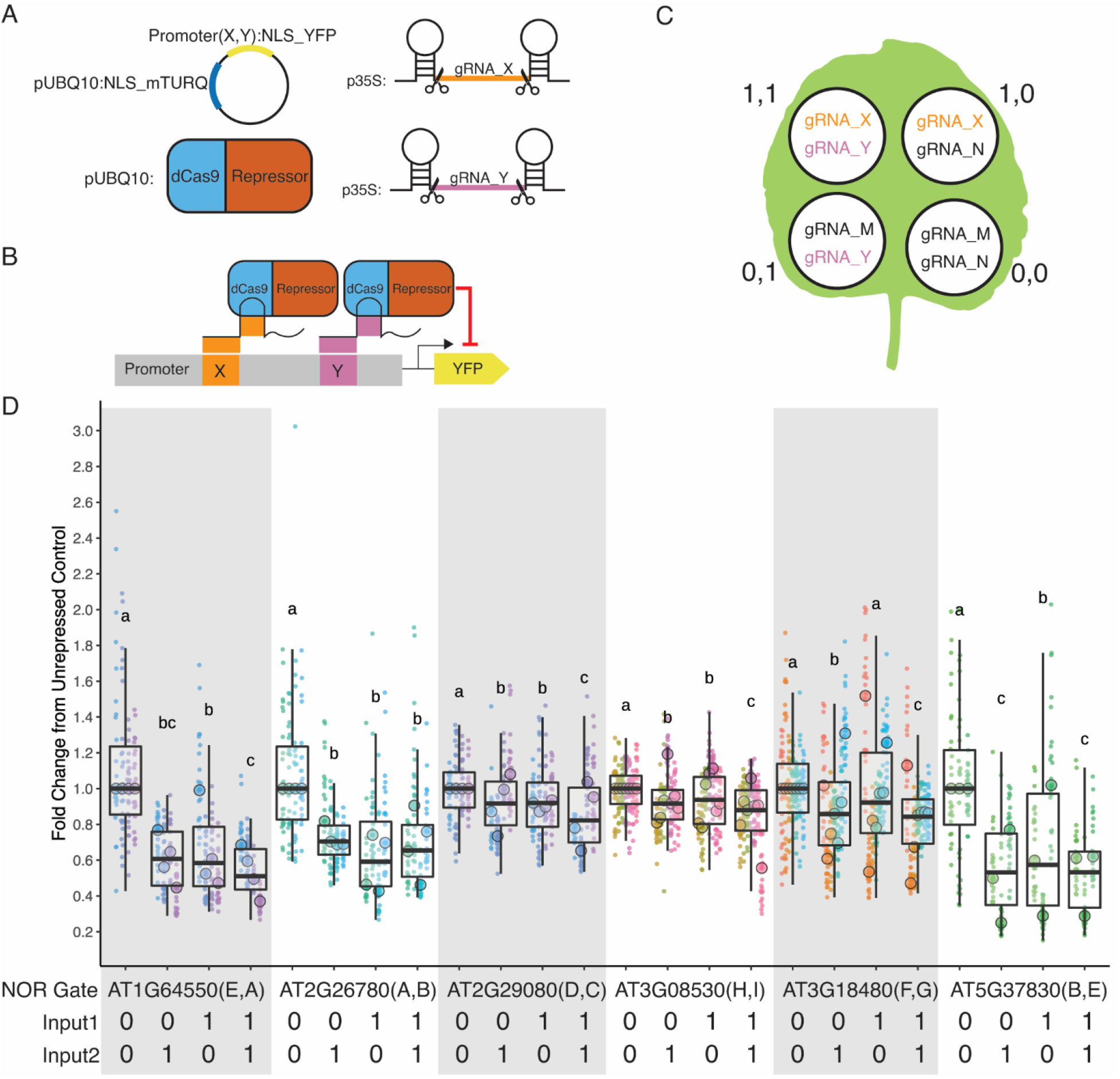
A) The four constructs co-injected for each injection. The injection always contains the mPromoter, dCas9-guided repressor, and the two self-cleaving input gRNAs. The gRNAs are denoted with X and Y representing a variable input. B) Schematic of the NOR gate when both input gRNAs are present. C) Pattern of injection for the four possible input combinations and the gRNAs used for each injection. (1,1) represents both guides are present while (0,0) represent neither are present. When a guide is not present, a non-matching gRNA is injected in its place, denoted here as gRNA_M or N. D) Five of the six mPromoters functioned as NOR gates. All guides apart from gRNA_F is independently repressible. Each biological replicate is represented by a beeswarm plot of individual datapoints collected from the plate reader as well as a single summarizing datapoint representing the median. The boxplot represents all biological replicates. The signal is measured as the YFP fluorescence (driven by the promoters being tested) divided by the mTURQ florescence (driven by pUBQ10). In each set of NOR gate injections, the (0,0) injection serves as the unrepressed control, and the dataset is normalized by dividing all values by the median of the unrepressed control on a per-leaf basis. The y-axis represents fold changes from the unrepressed control and each biological replicate of the control is centered on 1. Each color represents a unique leaf.

We first confirmed that the introduction of the target-sites did not abolish promoter expression (Figure2). While in most cases there was little difference in expression between modified and native promoters, in one case [64550(E,A)], the expression level increased dramatically, possibly due to introduction of new TF binding sites at the junction of the introduced gRNA target-site or disrupting a binding site for a repressor (Supplemental Figure 3). The repressibility of the modified promoters were tested in *N. benthamiana*, and five of the modified promoters (mPromoters) functioned as NOR gates while AT3G18480(F,G) acted as a NOT gate with input2 (gRNA_G) (Figure4D). Of the NOR gates, AT2G26780(A,B) was the best gate, with the promoter repressed to the same extent with either or both inputs. AT1G64550(E,A), AT2G29080(D,C) and AT3G08530(H,I) all displayed additive effects where having both inputs gave stronger repression than just having one alone. AT5G37830(B,E) had a well repressed target-site with input2 (gRNA_E) while input1 (gRNA_B) alone resulted in a weaker repressed state. The result displayed as normalized florescence and not as fold repression can be found in Supplemental Figure4.

## Discussion

Constitutive promoters are essential staples in stocking the synthetic biology toolbox. They are versatile due to their wide expression coverage, and form the foundation from which many synthetic promoters are built. Here, we report on the establishment of a pipeline to find the most stably expressing promoters in *Arabidopsis*. We successfully used this approach to identify sixteen promoters that are predicted to be more stably expressed than some of the most widely used native plant constitutive promoters, and showed they can drive expression in transient transformations of *N. benthamiana*. We attempted to capture the expression pattern of these promoters in stably transformed *Arabidopsis* using the visual RUBY reporter, and uncovered limitations in its utility. Lastly, we engineered repressible versions of six promoters and showed that five of these can function as NOR logic gates.

One of the biggest challenges in having a small selection of promoters to choose from is the need to repeat the use of the same promoter in larger constructs, which could pose challenges to stability. The promoters identified in this paper were selected from some of the most stably expressed genes available in the *Arabidopsis* genome and all have distinct sequences. A lack of promoter parts also means a lack of flexibility when it comes to the range of expression strength. Most of the promoters used in plant synthetic biology are quite strong and that is not ideal for every application. The availability of weaker constitutive promoters like those characterized here allows more flexibility in promoter choices when excess production of target proteins can be a problem. For example, they can be beneficial in avoiding toxic intermediates or optimizing flux in metabolic engineering projects [30,31].

The pipeline employed in this paper to arrive at new native constitutive promoters should be readily adaptable to other organisms, if there is sufficiently broad sampling of transcriptomes and a reference genome. The pipeline could also be modified to identify native promoters with particular expression patterns. One caveat is that the promoters that can be extracted in this way are, by definition, limited by what is naturally available in the organism. On the other hand, they have the advantage of already being assayed in a whole range of tissue types and developmental stages—a breadth of information that can be logistically challenging to collect for synthetic promoters. It will be interesting to see if synthetic devices made with these modified native promoters prove more resilient to mutation than those using fully engineered promoters, as these sequences have presumably maintained stable expression in the face of mutation and selection.

Working with native promoters also provided an opportunity to learn more about the biology of promoters themselves. Yamamoto and colleagues had suggested that plant promoters can be grouped into a few core promoter categories based on the presence or absence of certain location-sensitive motifs [32]. Interestingly, they reported that TATA-box containing promoters tend to be regulated promoters while Coreless promoters (promoters that don’t have any characteristic location-sensitive motifs) tend to be constitutively expressed. The vast majority of the constitutive promoters used today in plant synthetic biology are from the TATA promoter class, and we also have a much better understanding of how their expression is regulated [16]. If the goal is to find constitutive promoters, however, the analysis by Yamamoto and colleagues would suggest that we should look to Coreless promoters instead. Indeed, only 9% (3/33) of the candidate genes identified in this study contain TATA boxes, while 45% (15/33) are Coreless (Supplemental Table2).

The ability to selectively activate and repress genes provides the tools necessary to perform Boolean logic, which would allow more complex computations [33]. Plants naturally perform complex computations to determine when and where a gene should be expressed by integrating internal and external signals, and genetic logic gates provide a modular way to synthetically construct these input-output relationships by using simple genetic parts. There are many ways to achieve the different logic operations using molecular biology [30]. A NOR gate is powerful in that it can be used to construct any logic gate by just stringing together multiple NOR gates, and its efficacy had been demonstrated in yeast [34]. To date, the feasibility of building more complex logic circuits in plants has been hindered by the lack of unique and strongly repressible promoter parts. With just our design constraints and no additional refinement, five of the six NOR gates built showed the correct behavior. Additional design-build-test cycles in the future can help optimize the individual gRNA target sites and lower the overall OFF state, opening up a host of complex, synthetic plant logic operations in the future.

## Methods

### Downloading and processing RNA-seq datasets

We used a custom UseGalaxy pipeline to process the RNA-seq datasets [35]. SRR accession codes from BioProject IDs PRJNA314076 (138 samples; Klepikova et al., 2016), PRJNA268115 (20 samples; Klepikova et al. 2015), PRJNA324514 (32 samples), PRJNA194429 (2 samples; Loraine et al., 2013) were input into “Faster Download and Extract Reads in FASTQ (Galaxy Version 2.10.8+galaxy0)” with default settings. The FASTQ files were pipped into “FastQC (Galaxy Version 0.73+galaxy0)” and “Trimmomatic (Galaxy Version 0.38.0)” with sliding window trimming averaging across 4 bases with required average quality 20, and a minimum read length of 36. The trimmed files were input into “HISAT2 (Galaxy Version 2.1.0+galaxy5)” with reference genome assembly TAIR10 and Araport11 genome annotation from The *Arabidopsis* Information Resource (TAIR). Minimum intron length was set to 60, and maximum intron length was set to 6000 [36]. Features from the Araport11 annotation were counted with “htseq-count (Galaxy Version 0.9.1)” set to Union Mode and counting only reads within regions defined as “exons” in the Araport11 annotation while not counting non-unique/ambiguous reads [20]. The counted features were downloaded, and subsequent analysis was done in R [37].

### Identifying stable promoters

All samples excluding the stress dataset (PRJNA324514) were normalized together using the Median Ratios method from the DESeq2 package in R [38]. Coefficient of variation (CV) for each gene was calculated from the normalized data. Genes with the lowest 3% CV were kept for further analysis. Stress dataset from PRJNA324514 was normalized with “mature whole third leaf” from PRJNA314076 and its CV calculated on its own, separate from the rest of the data.

### Extracting promoter and terminator sequences

Promoter + 5’UTR region (from before the start codon and extending upstream till the first annotated neighboring gene or to a maximum of 2kb from the transcription start site, whichever is shorter) and 3’UTR + terminators (from after the stop codon and extending downstream till the first annotated neighboring gene or to a maximum of 250bp past the transcription end site, whichever is shorter) of the remaining genes were extracted using the Araport11 genome annotation and the “3000bp upstream and downstream” sequence files from the TAIR website. The extracted sequences were screened for BbsI and BsaI restriction enzyme cut sites and only those without were kept. Any genes with their promoter + 5’UTR and 3’UTR + terminator overlapping annotations from neighboring genes in the Araport11 annotation were also removed.

### Transcription factor binding site prediction

The promoter sequences of the remaining genes were uploaded onto PlantRegMap using the “Binding Site Prediction” function [27] and the predicted motifs for each promoter sequence were downloaded. Only genes that can fit two 23bp gRNA target sites at least 67bp apart without interrupting any of the predicted motifs while being within 500bp of the TSS were kept.

### Data release

The R analysis pipeline and the raw counts output from UseGalaxy can be found on Github (https://github.com/Nemhauserlab/StablePromoters) and in Supplemental Data1.

### Annotating candidate genes

For the final 33 candidate genes, CV for the Stress Dataset (StressCV) and promoter and terminator sequences were extracted as described above. The promoter core type was annotated from Tokizawa et al. 2017. A list of experimentally determined circadian genes in *Arabidopsis* was downloaded from CGDB [28], and any UniprotKB identifiers were converted to ATG identifiers with the Uniprot Retrieve/ID mapping tool. Gene Descriptions (Representative Gene Model Name, Gene Description, Gene Model Type, Primary Gene Symbol, and All Gene Symbols) were retrieved from TAIR.

### Construction of plasmids

Promoter + 5’UTR and 3’UTR + terminator for each candidate genes as defined above were cloned with PCR from extracted genomic Col-0 DNA into their respective MoClo level zero acceptors (pICH41295 and pICH41276 respectively) [25]. The promoter and terminator pair of the candidate genes were paired with nuclear localized Venus to make level one constructs in “position one”. Venus level one constructs were paired with pUBQ10 promoter driving nuclear localized mTURQ with an Act2 terminator from the MoClo Plant Parts Kit (pICH44300) in “position two” to form ratio-metric lvl2s with a binary Ti vector backbone (pAGM4673 or pAGM4723). RUBY from [29] was cloned into level zero constructs and then cloned directly into level-2 Ti vector backbone with the promoter and terminator pairs (pICH86966). List of primers and plasmid maps can be found in Supplemental Table 4 and 5 and Genbank files can be found in Supplemental Data 2.

### gRNA target-site introduction

Promoter regions within 500bp of the transcription start site were screened for predicted transcription factor (TF) binding sites with the PlantRegMap “Binding Site Prediction” webtool [27] as described above. gRNA target-sites were cloned into regions that does not disrupt any predicted TF binding sites through Gibson assembly [40]. Primers can be found in Supplemental Table 4.

### Agrobacterium infiltration

5mL cultures of *Agrobacterium* containing constructs to be injected were grown overnight at 30C with the appropriate antibiotics. A 25mL culture of P19 (Win and Kamoun 2004) was grown at the same time in the same conditions. On the following day, the overnights were centrifuged at 3000xg for 10 minutes. The pellets were resuspended with 1mL MMA (10 mM MgCl2, 10mM MES (pH 5.6), 100uM acetosyringone). The OD of the cultures were measured and about 1~2mL volume mixture with 5.0 OD for construct to be tested and 5.0 OD for P19 were prepared. The infiltration mix were rotated to mix at room temperature for 3hr before injecting into fully emerged *N. benthamiana* leaves with a 1mL syringe. The injections were always injected as triplicates on three separate leaves on three separate tobacco plants. Each leaf is also always injected a pUBQ10:mTURQ control.

### Fluorescence quantification in N. benthamiana

At 3 days post infiltration, the leaves were clipped off and visualized in the Azure C600 Westerblot imaging system with exposure times Cy5: 0 sec, Cy3: 15 sec, Cy2: 5 sec. Two hole-punches were taken out of representative regions of each injection, and damaged regions with high background fluorescence were avoided. The leaf discs were placed in a 96-well plate on top of 200uL of water, and the plates were read with a TECAN SPARK plate reader with YFP: excitation 506(15) and emission 541(15), Gain 100. mTurq: excitation 430(15) and emission 480(15), Gain 50. mScarlet: excitation 565(15) and emission 600(15), Gain 100. Settings: Multiple Reads Per Well; Circle (Filled) 4×4 with border 800uM. The output data was read into a custom R file for clean-up and visualization. Multiple biological replicates were assayed for each promoter being tested, and the three replicates closest to the median of all replicates were kept for visualization and statistical analysis. Each injection’s YFP value is subtracted by the median of the YFP value of the pUBQ10:mTURQ negative control on the same leaf, which serves as the background level of fluorescence. The YFP value is then divided by the mTURQ value for each injection to normalize across results. A Dunnett Test, a post hoc pairwise multiple comparison test from the DescTools package, was used to determine whether the injections were significantly different from the negative control [41]. The R code used can be found on Github (https://github.com/Nemhauserlab/StablePromoters) and in Supplemental Data1.

### Repression assays

Repression assays were performed using the modified promoters (mPromoters) with gRNA target-sites driving NLS-YFP and pUBQ10:NLS-mTURQ internal control as the reporter. The mPromoters (5OD) were co-injected with P19 (1OD), TPL repressor (1OD), self-cleaving gRNA_1 (1OD), and self-cleaving gRNA_2 (1OD). To test the mPromoters’ NOR gate functionality, two gRNA inputs were required. In cases where only one input is present, the other self-cleaving gRNA will be a non-matching guide to the mPromoter. When neither inputs are present, sometimes two non-matching gRNAs (1OD each) were co-injected and sometimes only one (2OD), but the final total OD were always consistent. The TPL repressor construct contains pUBQ1:tdTomato-pUBQ10:dCas9_TPL(N188) and is modified from Khakhar et al. 2018. Self-cleaving gRNAs were designed in accordance to Zhang et al. 2017, and the modifications (gRNA and complementary sequences) were introduced in one step using Q5 mutagenesis (NEB) and was placed in the MoClo pICH86988 acceptor with a 35S promoter. The four possible input combinations for the NOR gate for each promoter were always injected on the same leaf, and the result was read with a plate reader as described above. The YFP value of each injection was divided by the mTURQ value to normalize the data, and the value of each injection was divided by the median of the no-input control. An ANOVA followed by a Tukey’s Honest Significant Difference Test was used to determine significant differences in expression between samples. The R code used can be found on Github (https://github.com/Nemhauserlab/StablePromoters) and in Supplemental Data1. List of plasmid maps used can be found in Supplemental Table5 and Genbank files in Supplemental Data2.

### *RUBY expression in* Arabidopsis

Constructs with the candidate promoters driving RUBY were transformed into Col-0 through floral dipping method [43]. T1 seeds were selected on 0.5x LS + 50ug/mL Kanamycin + 0.8% bactoagar. Plates were stratified for 2 days, light pulsed for 6 hours then kept in the dark for 3 days. Resistant seedlings were transplanted to soil to collect T2 seeds. For each promoter lines, three representative T1 lines were chosen to have their T2 seedlings phenotyped, and for each line, 19 T2 seeds were plated on 120×120×17 square petri dishes with 0.5x LS + 0.8% bactoagar without selection. The plates were imaged on day 4, 8, and 12 post germination, and six representative seedlings were transplanted to soil. The plants were imaged with a digital camera on day 34. The flowers were imaged under a Leica S8AP0 dissecting scope. A representative leaf, a segment of the inflorescence, and silique were placed between two clear projector sheets and scanned with a flatbed scanner. List of plasmid maps used can be found in Supplemental Table5 and Genbank files in Supplemental Data2.

### qPCR

T2 seedlings were grown vertically on 0.5xLS + 0.8% Phytoagar and without selection. The plates were stratified at 4°C for 2 days. On day 12, approximately five seedlings per replicate were placed in 2mL tubes and powdered with a metal bead using a Retsch MM400 shaker after freezing the samples in liquid nitrogen. RNA was purified using an Illustra RNAspin Mini Kit (GE Healthcare). 1μg of extracted RNA was then used with the iScript cDNA synthesis kit (BIO-RAD). qPCR was performed using the iQ SYBR Green Supermix (BIO-RAD). PP2AA3 were used as a reference gene and the primers for PP2AA3 and RUBY can be found in Supplement Table1. The standard curves were established using a pool of all the cDNAs. Col-0 seedlings were also extracted on day 12 as four technical replicates. The qPCR were performed on a C1000 Thermal Cycler (BIO-RAD) and the result were read using the Bio-Rad CFX Maestro software and analyzed using standard methods [44].

## Supporting information

Supplemental_Table4_PrimerList

Supplemental_Table5_PlasmidList

Supplemental_Data1_Codes

Supplemental_Data2_Genbank

## Author Contributions

Experimental design and analysis by EJYY and JLN. Research performed by EJYY. Manuscript written by EJYY and JLN.

We thank Wesley George, Cassandra Maranas, Dr. Román Ramos Báez, Dr. Sarah Guiziou and Dr. Alexander Leydon for careful reading of the manuscript, as well as other members of the Nemhauser, Imaizumi, and Steinbrenner labs for their feedback on this project. We thank Dr. Nicholas J. Provart for his help with the RNA-seq datasets.

Information reported in the manuscript was previously presented at ICAR2022.

## Abbreviations

gRNA: guide RNA
TALE: Transcription activator-like effectors
CV: Coefficient of variation
NLS: Nuclear localization signal
OD: Optical Density

## Compliance with Ethics Guidelines

### Acknowledgements

This work was supported by the National Institute of Health (R01-GM107084), the National Science Foundation (IOS-1546873) and a Faculty Scholar Award from the Howard Hughes Medical Institute.

### Conflict of Interest

Eric J.Y. Yang and Jennifer L. Nemhauser declare that they have no conflict of interest

## Notes

### Competing Interest Statement

The authors have declared no competing interest.

