## Supplemental_Materials for "Expanding the synthetic biology toolbox with a library of constitutive and repressible promoters"

Supplemental Table1. qPCR primers for RUBY and PP2AA3

| Gene | Forward Primer | Reverse Primer |
|---|---|---|
| RUBY | AAACAGGGCAAGCTCGTGTA | ATCCGCAGTGGGTGAGAAAG |
| PP2AA3 | AACGTGGCCAAAATGATGC | AACCGCTTGGTCGACTATCG |

Supplemental Table2. Top 3% Candidate Genes

AGI is the *Arabidopsis* Genome Initiative locus code. Coefficient of variation (CV) and geometric mean (geom_mean) are calculated without the stress dataset while coefficient of variation for the stress dataset (StressCV) is given separately. The core promoter types (TATA, Ypatch, CA, GA, Coreless) are taken from Tokizawa et al. 2017, and “1” signifies that core type is predicted in the promoter. CGDB circadian genes are whether the promoter is found to be circadian regulated.

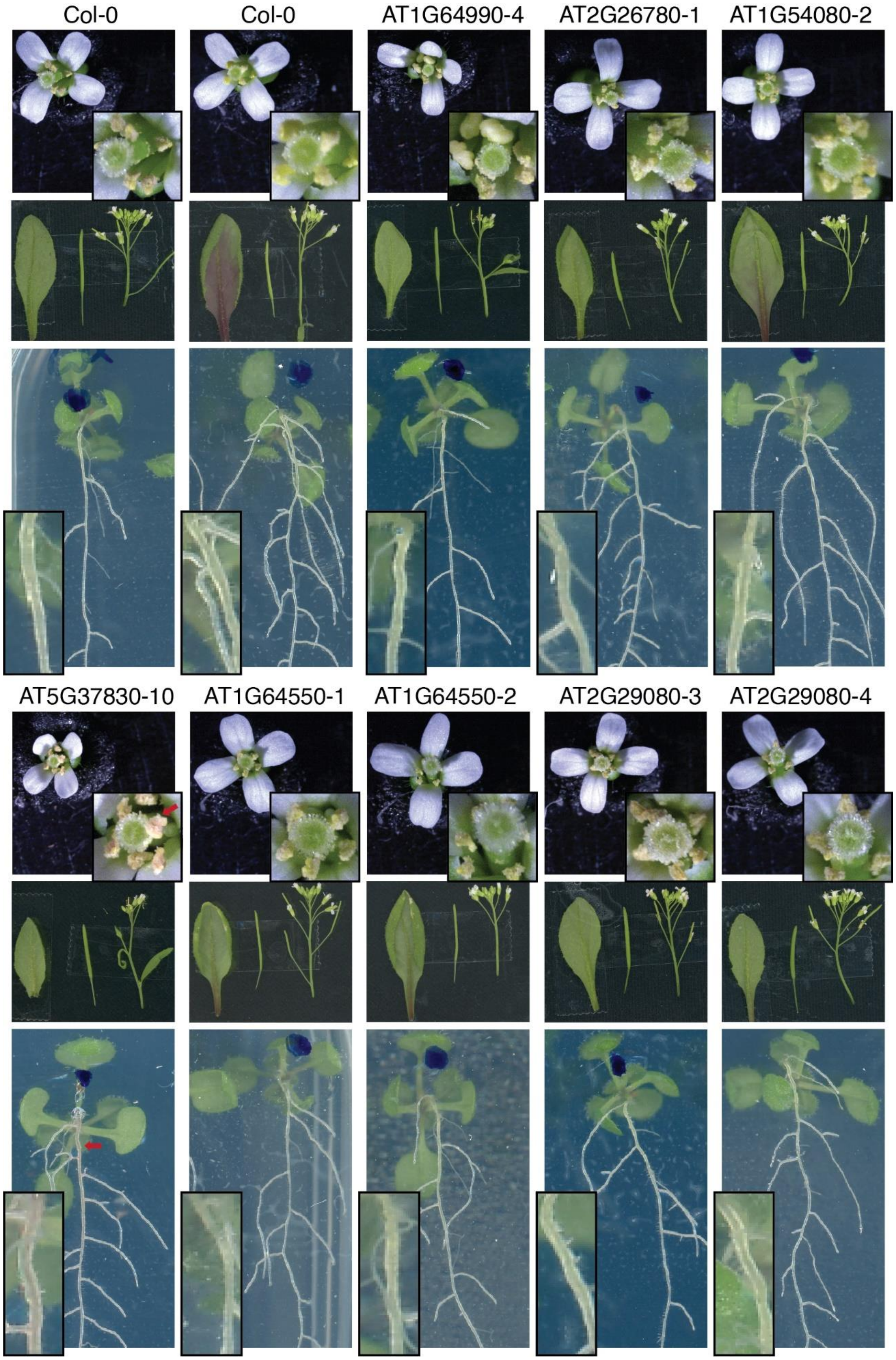
Col-0
Col-0
AT1G64990-4
AT2G26780-1
AT1G54080-2
AT5G37830-10
AT1G64550-1
AT1G64550-2
AT2G29080-3
AT2G29080-4

Supplemental Figure1. *Arabidopsis* T2 plants transformed with various promoters driving reporter RUBY. The flowers, siliques, and leaves are captured on day 34 while the seedling images are captured on day 12. The inset boxes are zoomed in pictures of their associated images. Areas where there are RUBY expression visible by eye is marked by the red arrow

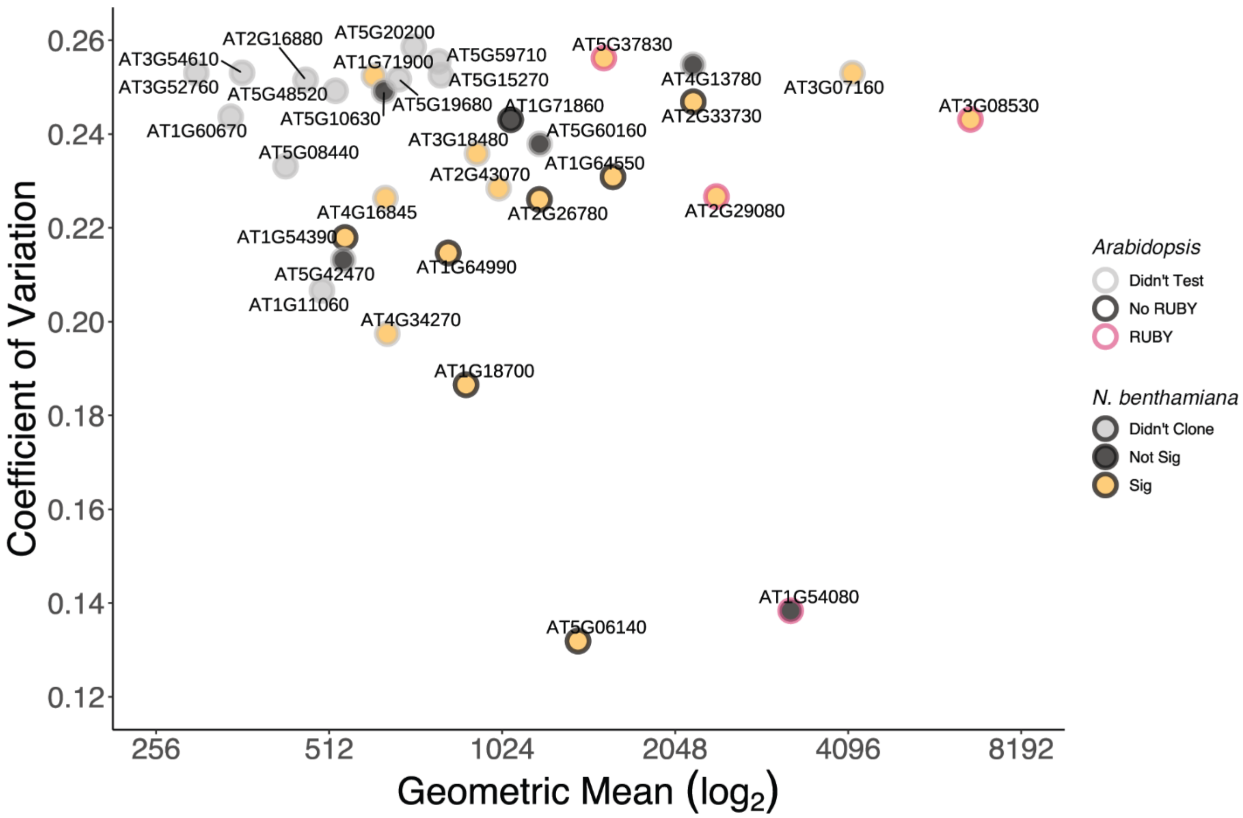


Supplemental Figure2. Final 33 candidates and the summary of experimental results. All final 33 candidates that passed through the pipeline were shown. Y-axis is the coefficient of variation, and the x-axis is the geometric mean on a log base-2 scale. The points were colored in grey if the construct wasn't cloned, black if the construct was cloned but showed no significant difference from negative control in transient *N. benthamiana* infiltration experiments, or yellow if the injection was significantly different from control. The points were outlined in grey if the RUBY construct wasn't tested in *Arabidopsis*, black if the promoter did not give visible RUBY expression, and red if at least one part of the tissue displayed visible RUBY expression.

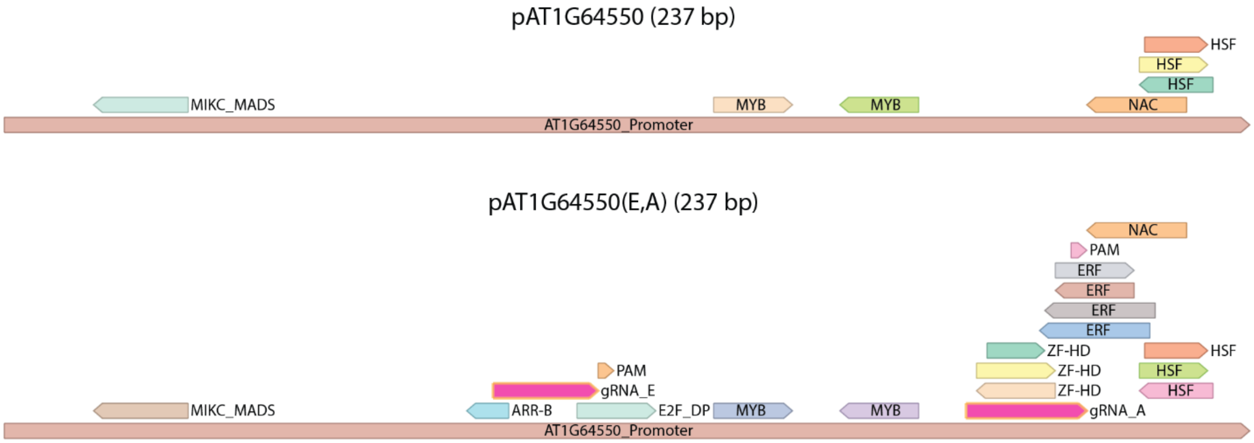


Supplemental Figure3. The introduction of gRNA_E and gRNA_A target-sites (highlighted in pink) in pAT1G64550 did not disrupt any predicted motifs but introduced additional ones.

Supplemental Table3. Guide Sequences

| gRNA | Guide Sequence + PAM | Source |
|---|---|---|
| A | GCAAAGGTGATTAACTGCAAAGG | [2] |
| B | AAAGGGGAAAAGAGTATTGGTGG | [3] |
| C | GGCAAGGCTGGCCAACCCATGGG | [3] |
| D | ACCCTGGCGGAGCTGATGGGTGG | [3] |
| E | TCTCAAGCTAGACTCTAGTGAGG | [3] |
| r3 | CATTGCCATACACCTTGAGGTGG | [4] |
| r5 | GAAGTCAGTTGACAGAGTCGTGG | [4] |
| r6 | GTGGTAACTTGCTCCATGTCTGG | [4] |
| r7 | CTTTACGTATAGGTTTAGAGTGG | [4] |

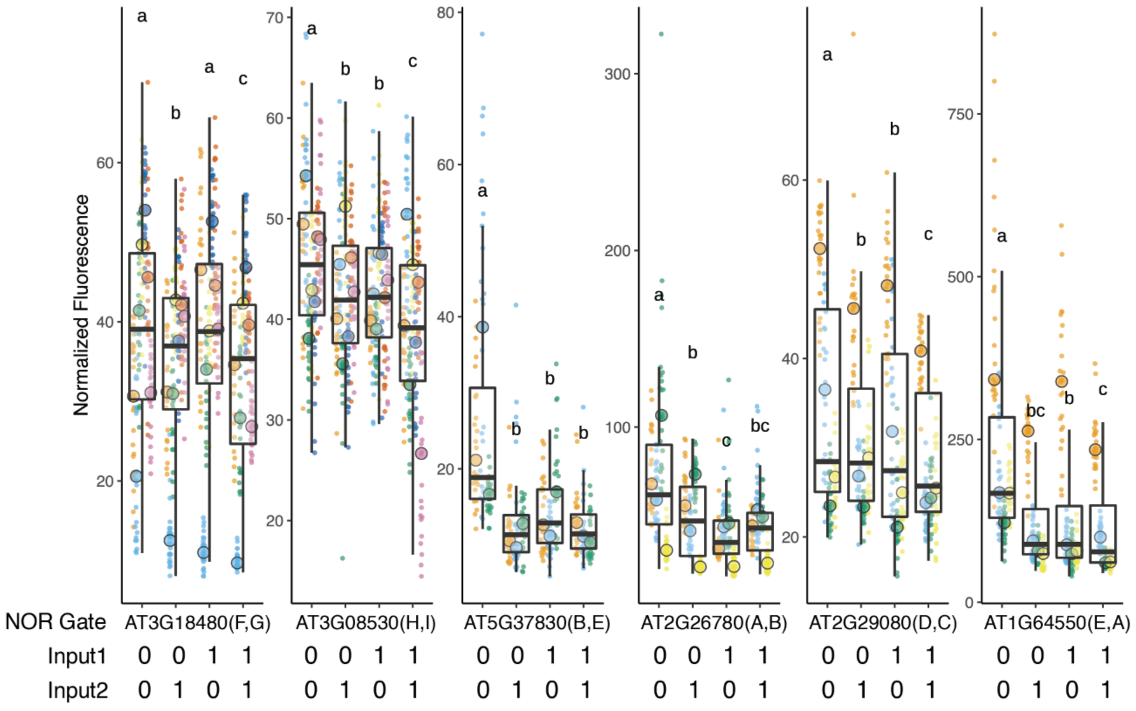


Supplemental Figure4. Normalized Repression Data. Each biological replicate represented by a beeswarm plot and their median is marked by the large circle. The boxplot represents all the replicates together. The y-axis is normalized fluorescence by having the mPromoter:NLS_YFP signal divided by pUBQ10:NLS_mTURQ signal within the same construct. Each input for a given condition can be either ON (1) or OFF (0), and each NOR gate can accept four possible combinations of the two inputs.

Supplemental Table4. List of primers used in the experiment

Supplemental Table5. List of plasmids used in the experiments, plasmid names correspond to Genbank files in Supplemental Data2.

Supplemental Data1. All the data files and codes used in the experiment

Supplemental Data2. All the plasmid maps of used in the experiment in Genbank format.
